# Enhancing t-SNE Performance for Biological Sequencing Data through Kernel Selection

**DOI:** 10.1101/2023.08.21.554138

**Authors:** Prakash Chourasia, Taslim Murad, Sarwan Ali, Murray Patterson

## Abstract

The genetic code for many different proteins can be found in biological sequencing data, which offers vital insight into the genetic evolution of viruses. While machine learning approaches are becoming increasingly popular for many “Big Data” situations, they have made little progress in comprehending the nature of such data. One such area is the t-distributed Stochastic Neighbour Embedding (t-SNE), a general-purpose approach used to represent high dimensional data in low dimensional (LD) space while preserving similarity between data points. Traditionally, the Gaussian kernel is used with t-SNE. However, since the Gaussian kernel is not data-dependent, it determines each local bandwidth based on one local point only. This makes it computationally expensive, hence limited in scalability. Moreover, it can misrepresent some structures in the data. An alternative is to use the isolation kernel, which is a data-dependent method. However, it has a single parameter to tune in computing the kernel. Although the isolation kernel yields better performance in terms of scalability and preserving the similarity in LD space, it may still not perform optimally in some cases. This paper presents a perspective on improving the performance of t-SNE and argues that kernel selection could impact this performance. We use 9 different kernels to evaluate their impact on the performance of t-SNE, using SARS-CoV-2 “spike” protein sequences. With three different embedding methods, we show that the cosine similarity kernel gives the best results and enhances the performance of t-SNE.

## 1 Introduction

In Machine Learning (ML), kernels are frequently used to solve challenges involving calculating object similarity between pairs of objects. The t-distributed Stochastic Neighbour Embedding (t-SNE) [15] is a popular solution used widely by researchers to reduce this dimensionality. Its goal is to project high-dimensional datasets into lower-dimensional spaces while maintaining data point similarities as indicated by the Kullback-Liebler (KL) divergence. This paper reviews the use of different kernels and their impact on t-SNE for SARS-Cov-2 ^1^ sequence data and its visualization in low dimension. We also assess these kernels in terms of classification and clustering. The vast global spread of COVID-19 spurred this, pushing viral sequence research into the “Big Data” sphere. This presents challenges since highly dimensional data cannot be used directly for machine learning solutions.

Since spike protein sequences cannot be used directly as input to machine learning (ML) models, we must first convert the sequences into a fixed-length numerical representation. For this purpose, using feature engineering-based methods is a popular choice, as proposed in many studies recently [12,2,10,1]. It has been shown that different embeddings can yield different results in terms of classification [3] and clustering [16,4,5,6] of SARS-CoV-2 spike sequences.

The t-SNE is a method to visualize high-dimensional data by mapping each point to a low-dimensional space (2 or 3 dimensions). It aims to preserve the structure of the data. In the literature, the Gaussian kernel is used by default for t-SNE-based visualization [25]. However, recent studies [25,8] show that the Gaussian kernel may not always be the best choice for t-SNE-based visualization, as it is computationally expensive and could perform worse than the isolation kernel (a data-dependent kernel). We argue in this paper that even using the isolation kernel [25] may not be the better option when dealing with biological sequences. We demonstrate that, when evaluating the t-SNE method both subjectively and objectively, numerous kernels outperform the Gaussian and isolation kernel using various embedding techniques(three feature engineering-based embeddings). In this paper, our contributions are the following:

1. We show that the cosine similarity-based kernel is a better choice for t-SNE as compared to 8 other kernel methods, including Gaussian and isolation kernels, in the case of SARS-CoV-2 spike sequences.
2. We evaluate the performance of the t-SNE model on both objective and subjective criteria, and report results for several kernel computation approaches.
3. We show that the cosine similarity kernel is better in terms of computational runtime and pairwise distance preservation in low dimensions as compared to the Gaussian and isolation kernel on SARS-CoV-2 sequences. Therefore, it could be a potential candidate for efficient t-SNE computation for the eventually larger sets of biological sequences [19].

The rest of the paper is organized as follows: Section 2 discusses the related work. Section 3 and Section 4, we discuss the methods used to compute the kernel matrix and detail for computing the t-SNE using the kernel matrix. Section 5 contains details of the different embeddings we use. Section 6 contains experimentation and data statistics. In Section 7, we evaluate the impact of different kernels on t-SNE. Finally, we conclude the paper in Section 8.

## 2 Related Work

Data visualization is an important task. Using t-SNE, originally introduced in [15], has made this task easy. Authors in [3,8] use t-SNE to visualize different variants in the coronavirus protein sequence data. It has also been found that clustering the COVID-19 protein sequences using k-means is also related to the patterns shown in the t-SNE plots [4,20,6,7].

Authors in [9] proposed Symmetric stochastic neighbor embedding (SNE) to get the 2D representation of the high dimensional data. An extension of t-SNE, called Heavy-tailed SNE is proposed in [24], which considers different embedding similarity functions. Authors in [22] propose a method, called t-Distributed Stochastic Triplet Embedding, based on the idea of similarity triplets to consider similar points and discard dissimilar points in the embeddings. Some efforts have been made previously to speed up the computation of t-SNE [21] using tree-based algorithms. Authors in [25] show that using the isolated kernel within the t-SNE could improve the visualization in 2D and its runtime compared to the Gaussian. A decentralized data stochastic neighbor embedding (dSNE) is proposed in [17], visualizing the decentralized data. The differentially private version of dSNE (DP-dSNE) is proposed in [18]. Although contemporary t-SNE methods are effective on well-known datasets like MNIST, it is unclear if those methods will be just as effective when applied to biological protein sequences.

## 3 Kernel Matrix Computation

Table 1 describes different methods we use to compute the kernel matrix.

**Table 1:**
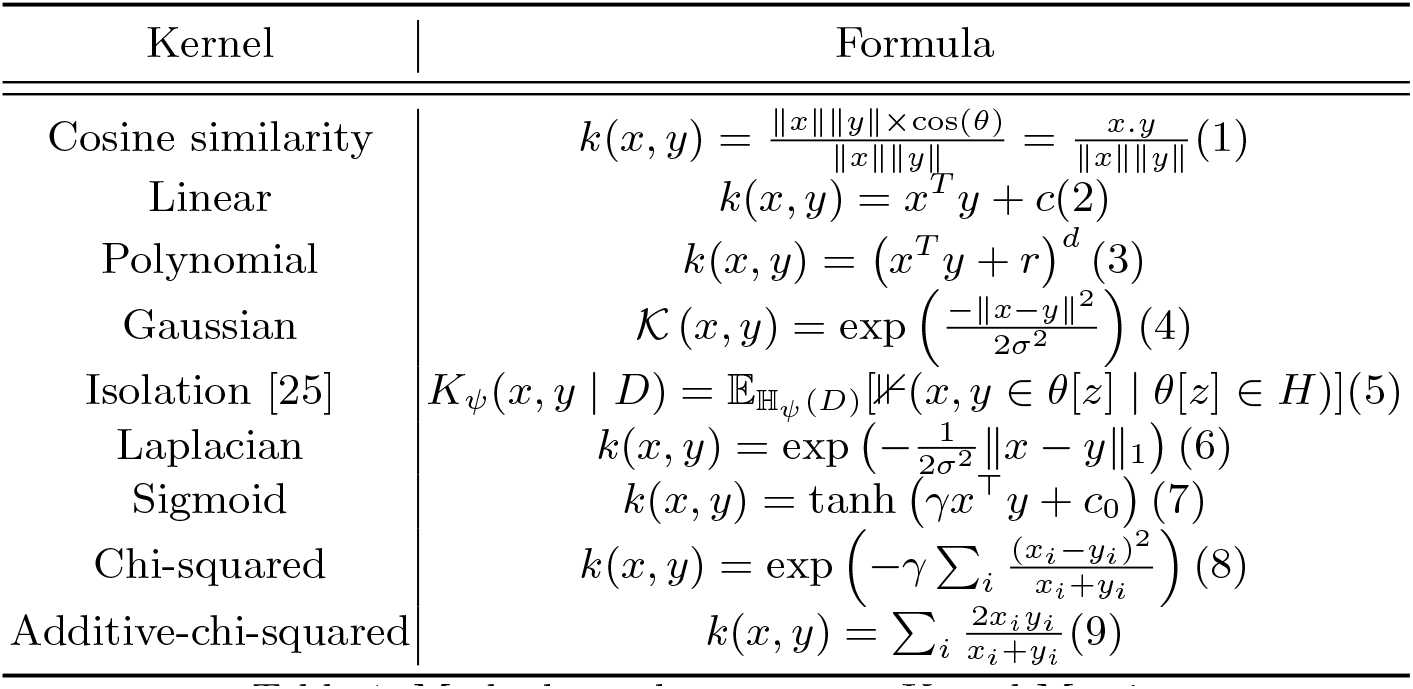
Methods used to compute Kernel Matrix.

## 4 Using Kernel Matrix for t-SNE

The t-SNE method takes an *n × n* kernel matrix as input, where *n* is the total number of sequences (embedding vectors), and produces a low dimensional representation *d*, where *d* = {1, 2, *· · ·, n* − 1}. More formally, given the high dimensional (HD) data, the idea of t-SNE is to represent the data in low dimensions (LD) while preserving the pairwise distance between the embedding vectors i.e to keep the distances between points in LD *Y* as close as to in HD *X*. The t-SNE approach works as follows:

### Compute Conditional Probability or Pairwise Affinities

The first step in t-SNE is to calculate the Euclidean distances of each point from all other points in high dimensions. This can be done using different kernel functions. Later this distance between data points is converted into conditional probabilities which are also known as pairwise affinities or similarity matrices.

### High Dimensional Probability Computation

The conditional probability can be gathered to give joint distribution on pairs of points. This is gathered into a symmetric matrix and returned joint probability can be written as given in Equation 10 (as also given in [23]):

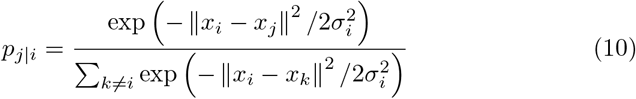

### Initial Solution Sampling

Initial solution *Y* is sampled with random initial values. These values are optimized to give the best lower dimensional representation of data points.

### Compute Low Dimensional Joint Probability

Similar to conditional probability in the high dimension, we compute it in the low dimension. Finally, gather these to get the low dimensional joint probability, which can be written as follows (also mentioned in [18]):

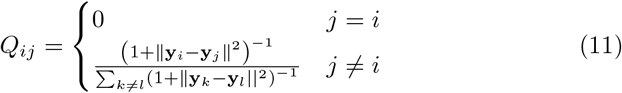

### Compute KL Divergence and Gradient

For these two distributions *P* in an HD and *Q* in an LD, to measure the distance between them we use KL divergence. It is used to find the variation or distribution among the distances in the data points. KL divergence is computed as:

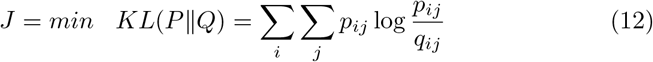

where *J* is the cost function, *P* and *Q* represents two different dimensions, and *y*_*i*_, *y*_*j*_ are points in *Q*. Take the derivative and calculate gradient descent Equation 13 (as mentioned in [23]) to get the minimum from *J*

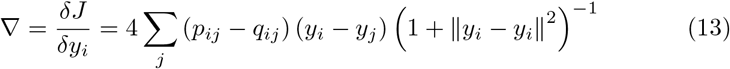

and keep updating *Y* for minimum *J* Equation 14. We apply t-distribution in this Stochastic Neighborhood Embedding (SNE), applying t-distribution on *Q* low dimension gives us a longer tail to give better visualization.

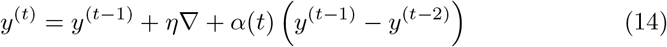

where *t T* represents t-time iterations.

In summary, the kernel function plays an important role here to give distances between the points in the original data. The workflow of t-SNE computation using the kernel matrix as input is illustrated in Algorithm 1.

#### Algorithm 1 t-SNE Computation.

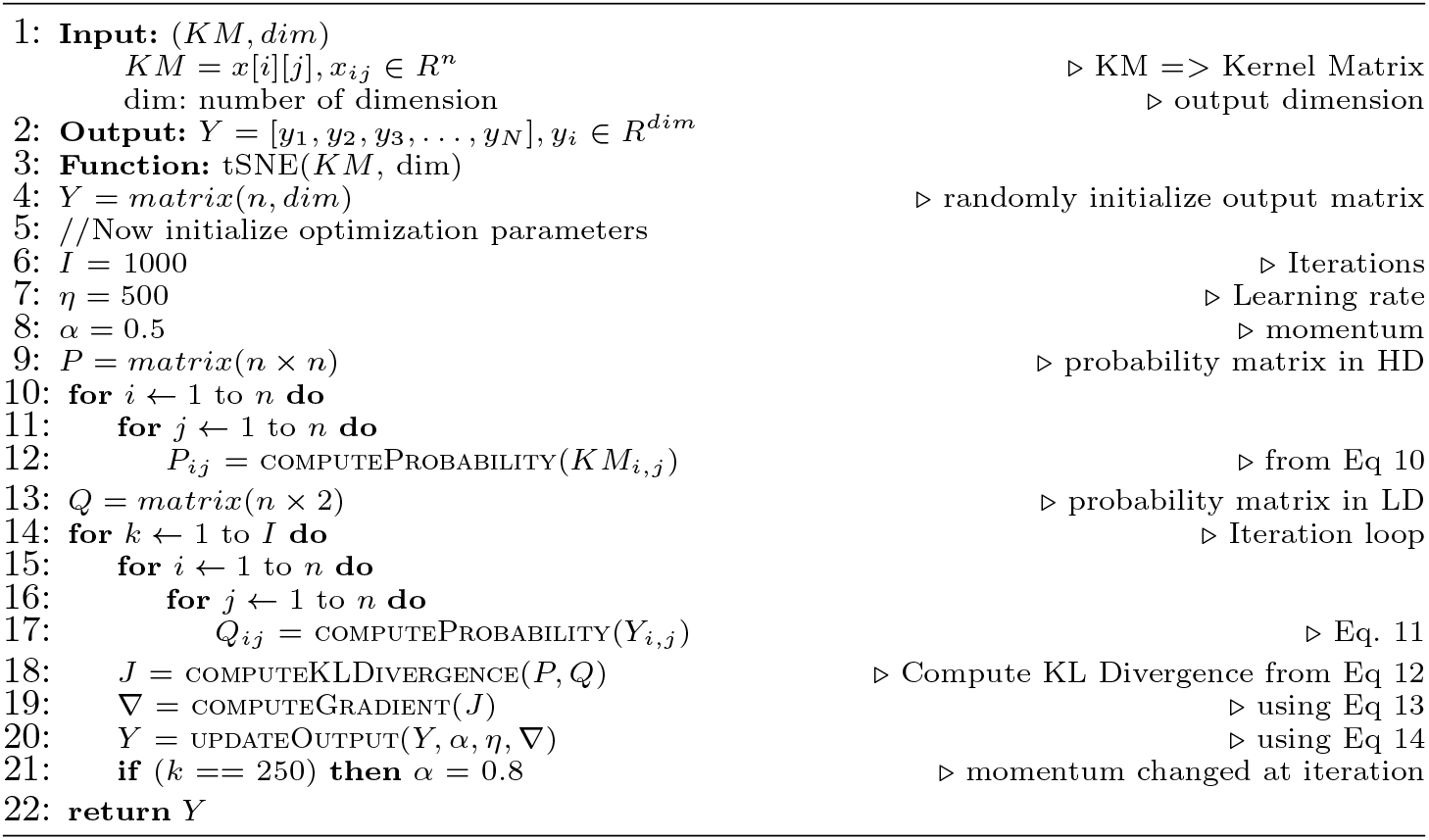

## 5 Feature Embeddings Generation

This section describes the three embedding methods we use to convert the biological sequences into a fixed-length numerical representation.

1. One Hot Encoding (OHE) - To convert the amino acids into numerical representation we use OHE [12,3].
2. Spike2Vec [2] - It generates a fixed-length numerical representations using the concept of *k*-mers (also called n-gram). It uses the sliding window concept to generate substrings (called mers) of length *k* (size of the window).
3. Minimizers - Using the *k*mer, the minimizer is computed as a substring (mer) of length *m* (where *m < k*) within that *k*mer. It is a lexicographical minimum in the forward and reverse order of the *k*-mer.

## 6 Experimental Setup

In this section, we discuss the dataset statistics followed by the goodness metrics used to evaluate the performance of t-SNE. All experiments are performed on Intel (R) Core i5 system with a 2.40 GHz processor and 32 GB memory.

### 6.1 Data Statistics

The dataset we use, we named Spike7k consists of sequences of the SARS-CoV-2 virus and is taken from well known database GISAID [11]. It has 22 unique lineages as the label with the following distribution: B.1.1.7 (3369), B.1.617.2 (875), AY.4 (593), B.1.2 (333), B.1 (292), B.1.177 (243), P.1 (194), B.1.1 (163), B.1.429 (107), B.1.526 (104), AY.12 (101), B.1.160 (92), B.1.351 (81), B.1.427 (65), B.1.1.214 (64), B.1.1.519 (56), D.2 (55), B.1.221 (52), B.1.177.21 (47), B.1.258 (46), B.1.243 (36), R.1 (32).

### 6.2. Evaluating t-SNE

For objective evaluation of the t-SNE model, we use a method called *k*-ary neighborhood agreement (*k*-ANA) method [25]. The *k*-ANA method (for different *k* nearest neighbors) checks the neighborhood agreement (*Q*) between HD and LD and takes the intersection on the numbers of neighbors. More formally:

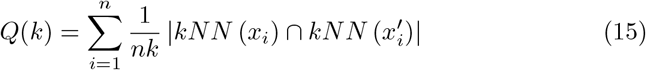

where *k*NN(*x*) is set of *k* nearest neighbours of *x* in high-dimensional and *k*NN(*x*^*′*^) is set of *k* nearest neighbours of *x* in corresponding low-dimensional. We use a quality assessment tool that quantifies the neighborhood preservation and is denoted by R(*k*), which uses Equation 15 to evaluate on scalar metric whether neighbors are preserved[13] in low dimensions. More formally:

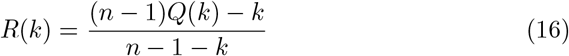

R(*k*) represents the measurement for *k*-ary neighborhood agreement. Its value lies between 0 and 1, the higher score represents better preservation of the neighborhood in LD space. In our experiment, we computed *R*(*k*) for *k* ∈ 1, 2, 3, …, 99 then considered the area under the curve (AUC) formed by *k* and R(*k*). Finally, to aggregate the performance for different k-ANN, we calculate the area under the *R*(*k*) curve in the log plot (*AUC*_*RNX*_) [14]. More formally:

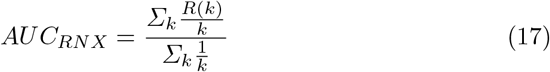

where *AUC*_*RNX*_ denotes the average quality weight for *k* nearest neighbors.

## 7 Subjective and Objective Evaluation of t-SNE

This section discusses the performance of t-SNE in subjective (using 2D scatter plots) and objective (using *AUC*_*RNX*_) ways for different embeddings and kernels.

### 7.1 Subjective Evaluation

To visually evaluate the performance of t-SNE, we use different embedding methods and plot the 2D visual representation to analyze the overall structure of the data. Figure 1 shows the top 2 performing kernels in terms of *AUC*_*RNX*_ score for respective embedding. Similarly, Figure 2 shows the worse 2 performing kernels.

**Fig. 1:**
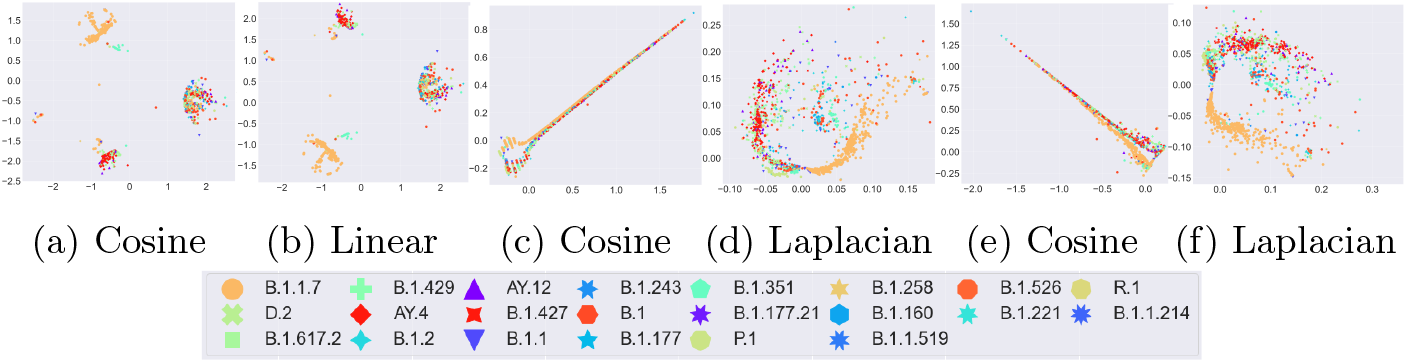
t-SNE plots for top-performing kernel methods for different embedding. This figure is best seen in color. (a), (b) is from OHE. (c), (d) are from Spike2Vec. and (e), (f) are from Minimizer encoding.

**Fig. 2:**
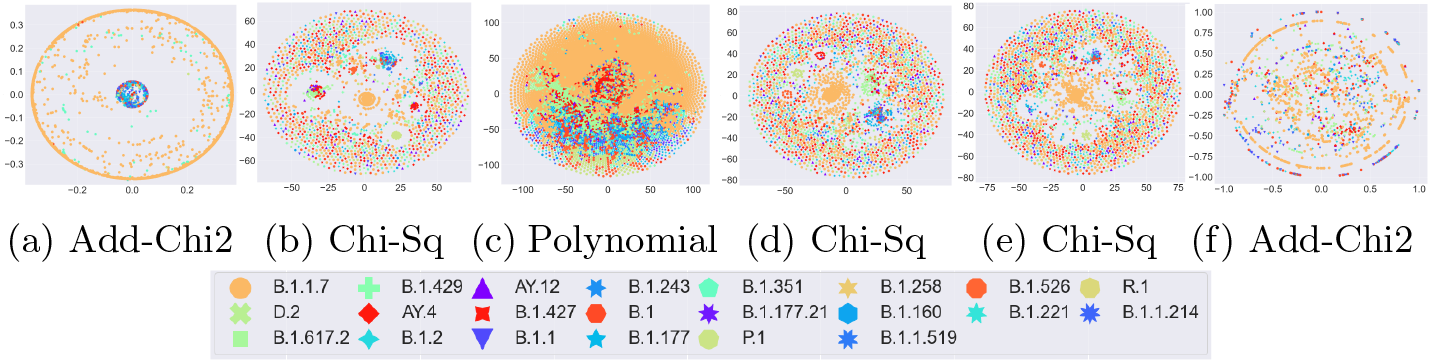
t-SNE plots for worst-performing kernel methods for different embedding. This figure is best seen in color. (a), (b) is from OHE. (c), (d) are from Spike2Vec. and (e), (f) are from Minimizer encoding.

**Fig. 3:**
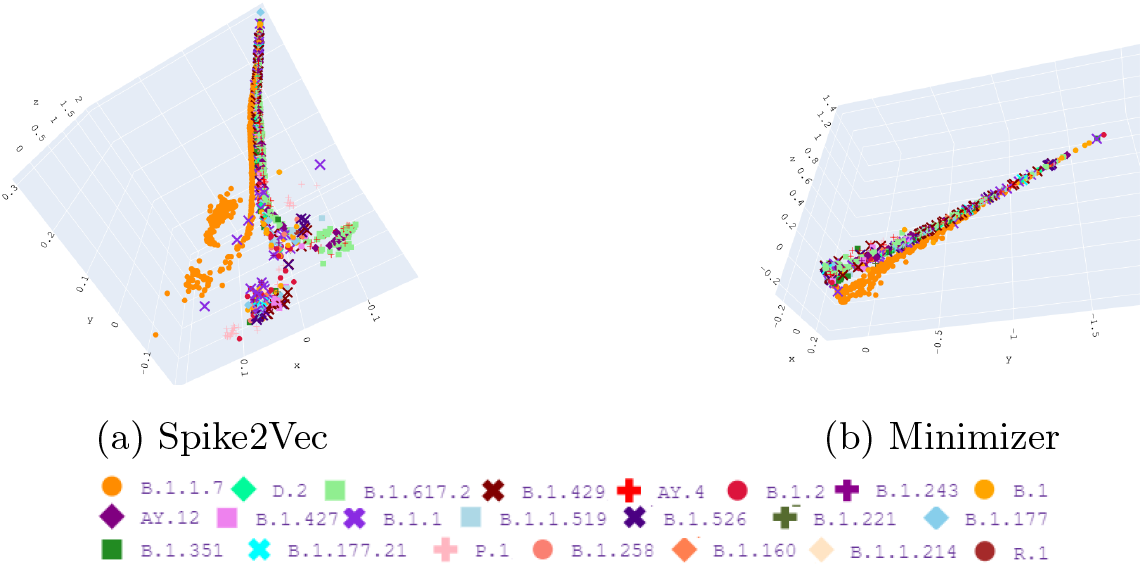
t-SNE 3d plots using Cosine similarity kernel.

#### OHE [12]

We analyzed the t-SNE plots for one-hot embedding for different kernel methods. For the Alpha variant (B.1.1.7), we can see that Gaussian, Isolation, Linear, and Cosine kernel can generate a clear grouping. However, for the other variants having a small representation in the dataset (e.g. B.1.617.2 and AY.4), we can see the Cosine and linear kernels are better than the Gaussian and Isolation. This could mean that the Gaussian and Isolation tend to be biased toward the more representative class (alpha variant).

#### Spike2Vec [2]

The t-SNE plots for Spike2vec-based embeddings using different kernel methods are evaluated. It is similar to OHE, where the Gaussian and isolation kernels group the alpha (B.1.1.7) variant almost perfectly, however, all other variants are scattered around in the plot.

#### Minimizer

Similarly in the t-SNE plots for minimizer embedding for different kernel methods for the Gaussian and isolation kernel, we can observe similar behavior for the alpha (B.1.1.7) variant.

We also show the 3D plots for t-SNE using the Cosine similarity kernel in Figure 3 for Spike2Vec and Minimizers-based embedding. We can see for Spike2Vec, the Alpha variant shows clear grouping. Similarly, the delta and epsilon variant also contains a few small groups.

### 7.2 Objective evaluation of t-SNE

The objective evaluation of tSNE is done using Equation 17. The goodness of t-SNE using kernel computation runtime, and *AUC*_*RNX*_ for different embedding approaches are reported in Table 2. An interesting insight, which we can observe is that the Cosine similarity-based kernel outperforms all other kernel methods in terms of kernel computational runtime and *AUC*_*RNX*_ value for all embedding methods. This means that not only the Cosine similarity-based kernel could be scaled easily on a bigger dataset, but also that its neighborhood agreement in high dimensional data vs. low dimensional data is highest. Therefore, we can conclude that for the biological protein sequences, using the cosine similarity kernel is better than the Gaussian or isolation kernel (authors in [25] argue that using the isolation kernel with t-SNE is better).

**Table 2:** *AUC*_*RNX*_ values for t-SNE using different kernel and encoding methods on Spike7k datasets. The kernel is sorted in descending order with the best at the top and the best values are shown in underlined and bold.

| OHE |  |  | Spike2Vec |  |  | Minimizer |  |  |
| --- | --- | --- | --- | --- | --- | --- | --- | --- |
| Kernel | Kernel<br>comp. time | $AUC_{RNX}$ | Kernel | Kernel<br>comp. time | $AUC_{RNX}$ | Kernel | Kernel<br>comp. time | $AUC_{RNX}$ |
| Cosine similarity | **<u>37.351</u>** | **<u>0.262</u>** | Cosine similarity | **<u>15.865</u>** | **<u>0.331</u>** | Cosine similarity | **<u>17.912</u>** | **<u>0.278</u>** |
| Linear | 51.827 | 0.260 | Laplacian | 721.834 | 0.260 | Laplacian | 983.26 | 0.242 |
| Gaussian | 94.784 | 0.199 | Gaussian | 54.192 | 0.235 | Sigmoid | 27.813 | 0.229 |
| Sigmoid | 57.298 | 0.190 | Linear | 30.978 | 0.197 | Gaussian | 65.096 | 0.206 |
| Laplacian | 2250.30 | 0.184 | Isolation | 24.162 | 0.189 | Isolation | 27.266 | 0.172 |
| Polynomial | 57.526 | 0.177 | Sigmoid | 30.037 | 0.168 | Polynomial | 28.666 | 0.166 |
| Isolation | 39.329 | 0.161 | Additive-chi2 | 495.221 | 0.121 | Linear | 27.503 | 0.165 |
| Chi-squared | 1644.86 | 0.131 | Chi-squared | 495.264 | 0.104 | Additive-chi2 | 576.92 | 0.125 |
| Additive-chi2 | 1882.45 | 0.110 | Polynomial | 28.427 | 0.059 | Chi-squared | 921.63 | 0.095 |

### 7.3 Runtime Analysis

The runtime for computing different kernel matrices with an increasing number of sequences is shown in Figure 4a and its zoomed version in Figure 4b. We can see the Cosine similarity outperforms all other kernel methods. The Gaussian takes the longest. Moreover, we can observe that the overall runtime increasing trend for most kernels is linear. We report the t-SNE computation runtime for the Cosine similarity with an increasing number of sequences in Figure 4c. We can see a linear increase in the runtime as we increase the sequences.

**Fig. 4:**
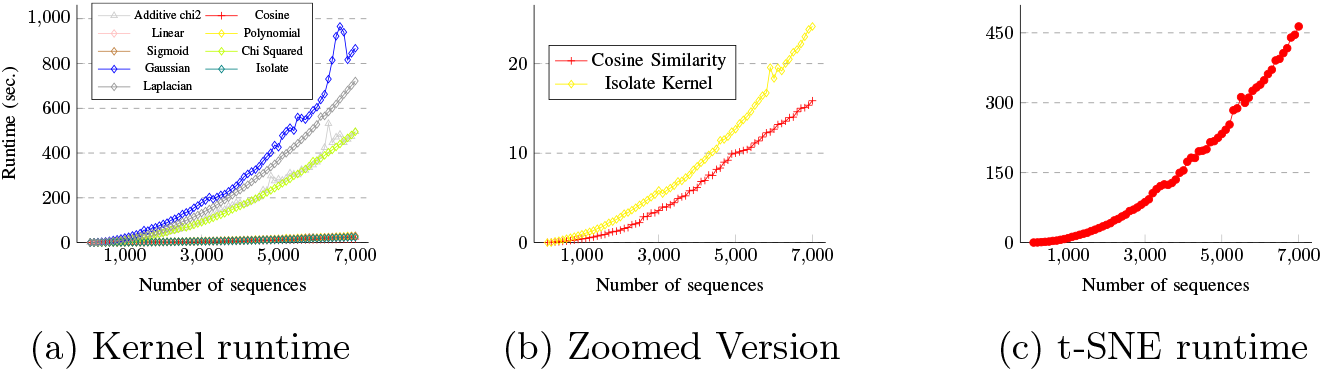
(a) and (b) shows the kernel computation Runtime for an increasing number of sequences (using Spike2Vec-based embedding). (c) shows the t-SNE computation runtime with an increasing number of sequences (using Spike2Vec-based embedding). This figure is best seen in color.

## 8 Conclusion

In this paper, we modify the original tSNE algorithm and show its performance using different kernel methods. We demonstrate that the cosine similarity-based kernel performs best among 9 kernels for t-SNE-based visualization. We show that, rather than using Gaussian or isolation kernel (as argued in previous works), the cosine similarity kernel not only yields better computational runtimes (hence better scalability) but also improves the performance of t-SNE. In the future, we will explore other biological data to evaluate the performance of the reported kernels. We also plan to use different embedding and kernel methods to explore the impact on the classification and clustering results for the biological sequences.

## Footnotes

1 The SARS-CoV-2 virus is the cause of the global COVID-19 pandemic.

